# SARS-CoV-2 infection activates dendritic cells via cytosolic receptors rather than extracellular TLRs

**DOI:** 10.1101/2021.09.02.458667

**Authors:** Lieve E.H. van der Donk, Julia Eder, John L. van Hamme, Philip J.M. Brouwer, Mitch Brinkkemper, Ad C. van Nuenen, Marit J. van Gils, Rogier W. Sanders, Neeltje A. Kootstra, Marta Bermejo-Jambrina, Teunis B.H. Geijtenbeek

## Abstract

Severe acute respiratory syndrome coronavirus 2 (SARS-CoV-2) causes coronavirus disease 2019 (COVID-19), an infectious disease characterized by strong induction of inflammatory cytokines, progressive lung inflammation and potentially multi-organ dysfunction. It remains unclear whether SARS-CoV-2 is sensed by pattern recognition receptors (PRRs) leading to immune activation. Several studies suggest that the Spike (S) protein of SARS-CoV-2 might interact with Toll-like receptor 4 (TLR4) and thereby activate immunity. Here we have investigated the role of TLR4 in SARS-CoV-2 infection and immunity. Neither exposure of isolated S protein, SARS-CoV-2 pseudovirus nor a primary SARS-CoV-2 isolate induced TLR4 activation in a TLR4-expressing cell line. Human monocyte-derived dendritic cells (DCs) express TLR4 but not ACE2, and DCs were not infected by a primary SARS-CoV-2 isolate. Notably, neither S protein nor the primary SARS-CoV-2 isolate induced DC maturation or cytokines, indicating that both S protein and SARS-CoV-2 virus particles do not trigger extracellular TLRs, including TLR4. Ectopic expression of ACE2 in DCs led to efficient infection by SARS-CoV-2. Strikingly, infection of ACE2-positive DCs induced type I IFN and cytokine responses, which was inhibited by antibodies against ACE2. These data strongly suggest that not extracellular TLRs but intracellular viral sensors are key players in sensing SARS-CoV-2. These data imply that SARS-CoV-2 escapes direct sensing by TLRs, which might underlie the lack of efficient immunity to SARS-CoV-2 early during infection.

**Author summary:** The immune system needs to recognize pathogens such as SARS-CoV-2 to initiate antiviral immunity. Dendritic cells (DCs) are crucial for inducing antiviral immunity and are therefore equipped with both extracellular and intracellular pattern recognition receptors to sense pathogens. However, it is unknown if and how SARS-CoV-2 activates DCs. Recent research suggests that SARS-CoV-2 is sensed by extracellular Toll-like receptor 4 (TLR4). We have previously shown that DCs do not express ACE2, and are therefore not infected by SARS-CoV-2. Here we show that DCs do not become activated by exposure to viral Spike proteins or SARS-CoV-2 virus particles. These findings suggest that TLR4 and other extracellular TLRs do not sense SARS-CoV-2. Next, we expressed ACE2 in DCs and SARS-CoV-2 efficiently infected these ACE2-positive DCs. Notably, infection of ACE2-positive DCs induced an antiviral immune response. Thus, our study suggests that infection of DCs is required for induction of immunity, and thus that intracellular viral sensors rather than extracellular TLRs are important in sensing SARS-CoV-2. Lack of sensing by extracellular TLRs might be an escape mechanism of SARS-CoV-2 and could contribute to the aberrant immune responses observed during COVID-19.

## Introduction

Severe acute respiratory syndrome coronavirus 2 (SARS-CoV-2) is a novel coronavirus that causes coronavirus disease 2019 (COVID-19)(1). COVID-19 emerged in 2019 in Wuhan, China(2), and has since spread globally causing a pandemic. The symptoms of COVID-19 vary amongst individuals, ranging from mild respiratory symptoms to severe lung injury, multi-organ dysfunction and death(3–6). Increasing evidence suggests that disease severity depends not solely on viral infection, but also on an excessive host pro-inflammatory response, whereby high concentrations of pro-inflammatory cytokines result in an unfavorable immune response and induce tissue damage(7, 8). The events leading to excessive pro-inflammatory responses are not completely understood. Therefore, it is necessary to elucidate the mechanisms that are triggered by SARS-CoV-2 to induce innate and adaptive immune responses.

Innate immune cells express pattern recognition receptors (PRRs) that recognize pathogen-associated molecular patterns (PAMPs) and subsequently orchestrate an immune response against pathogens(9). Dendritic cells (DCs) are essential immune cells that function as a bridge between innate and adaptive immunity. DCs express various PRR families such as Toll-like receptors (TLRs) and cytosolic RIG-I-like receptors (RLRs) that are triggered upon virus interaction or infection(10). DCs are therefore essential during SARS-CoV-2 infection to sense infection and instruct T and B cells for efficient antiviral immune responses. However, it is unclear whether and how SARS-CoV-2 is sensed by DCs.

SARS-CoV-2 Spike (S) protein uses angiotensin converting enzyme 2 (ACE2)(11, 12) as receptor for infection. However, besides interacting with ACE2, recent *in silico* analyses suggest that the Spike (S) protein could also potentially interact with members of the TLR family, in particular TLR4(13, 14). TLR4 is abundantly expressed on DCs(15, 16), and therefore TLR4 signaling could be involved in induction of pro-inflammatory mediators. Other studies using cell lines and SARS-CoV-2 S protein support a potential interaction of TLR4 with the S protein(17–19). However, it remains unclear whether infectious SARS-CoV-2 virus is sensed by TLR4 and whether this interaction induces DC activation and initiation of immunity.

Here, we have investigated how SARS-CoV-2 is sensed by human DCs. Neither recombinant S protein, SARS-CoV-2 pseudovirus nor a primary SARS-CoV-2 isolate induced immunity in TLR4-expressing cell lines or DCs, indicating that TLR4 or other extracellular TLRs are not involved in SARS-CoV-2 infection. However, ectopic expression of ACE2 on DCs led to infection by SARS-CoV-2 and induction of type I interferon (IFN) and cytokines. These data imply that intracellular PRRs rather than transmembrane TLRs are involved in instigating an immune response against SARS-CoV-2.

## Results

### SARS-CoV-2 S protein does not trigger TLR4

To assess whether TLR4 acts as a sensor of S protein of SARS-CoV-2, we treated a TLR4-expressing HEK293 cell line (293/TLR4) with SARS-CoV-2 recombinant S protein or S nanoparticles(20) and determined activation by measuring interleukin (IL)-8. Neither S protein nor S nanoparticles induced IL-8 secretion by 293/TLR4 cells, in contrast to the positive control LPS (Fig 1A). The parental 293 cells did not induce IL-8 upon treatment with S protein or S nanoparticle and LPS. These data suggest that S protein of SARS-CoV-2 does not trigger TLR4.

**Fig 1:**
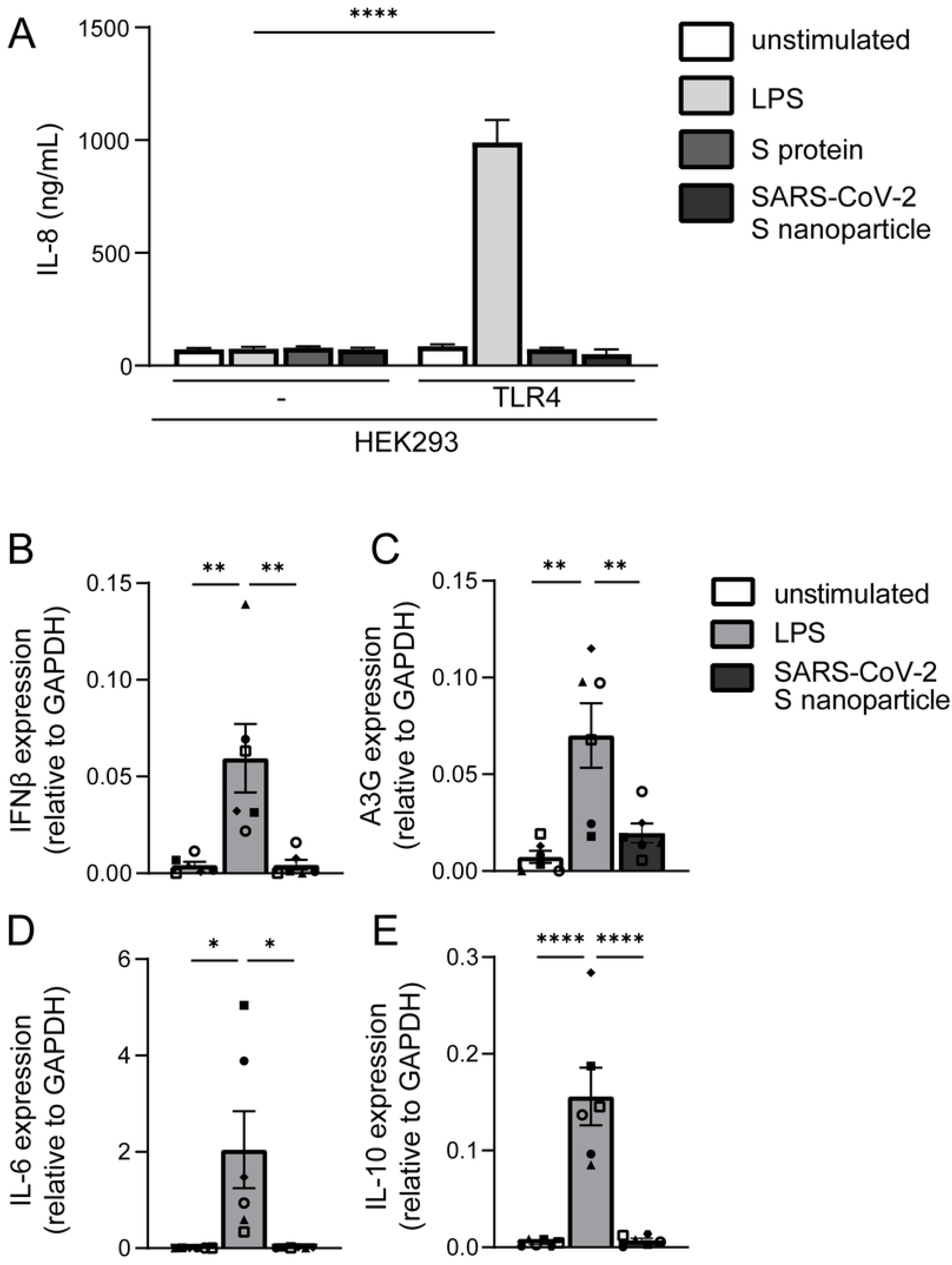
S protein and SARS-CoV-2 S nanoparticle do not trigger TLR4. (A) 293 cells or 293/TLR4 cells were exposed to LPS, SARS-CoV-2 S protein or S nanoparticles for 24h. IL-8 production was determined by ELISA. (B-E) Primary dendritic cells were exposed to LPS or SARS-CoV-2 S nanoparticles for 8h. Expression of IFNβ (B), A3G (C), IL-6 (D) and IL-10 (E) was determined with qPCR. Data show the mean values and SEM. Statistical analysis was performed using (A) two-way ANOVA with Šidák’s multiple comparisons test, or (B-E) one-way ANOVA with Tukey’s multiple comparisons test. (A) ****p<0.0001 (n=3). (B-E) ***p<0.001; **p<0.01; *p<0.05 (n=6).

Primary monocyte-derived DCs express TLR4 but also other TLRs(21). We therefore exposed primary human DCs to SARS-CoV-2 S nanoparticles and assessed cytokine production by qPCR. Treatment of DCs with S nanoparticles did neither induce type I interferon (IFN) nor cytokines. (Fig 1B-E). The positive control LPS induced IFNβ (Fig 1B) and the interferon-stimulated gene (ISG) APOBEC3G (A3G) (Fig 1C) as well as cytokines IL-6 and IL-10 (Fig 1D, E). These data strongly suggest that S protein from SARS-CoV-2 does not trigger extracellular TLRs on DCs.

### SARS-CoV-2 virus particles do not trigger TLR4

To assess whether TLR4 plays a role in SARS-CoV-2 entry and replication, we ectopically expressed ACE2 on 293 and 293/TLR4 cell lines and infected the cells with SARS-CoV-2 pseudovirus that expresses the full-length S glycoprotein from SARS-CoV-2 and contains a luciferase reporter gene(22). Infection was determined by measuring luciferase activity. SARS-CoV-2 pseudovirus infected ACE2-positive 293 and 293/TLR4 cells but not the parental 293 and 293/TLR4 cells (Fig 2A). TLR4 expression did not affect infection, as infection was comparable between 293/ACE2 and 293/TLR4/ACE2 cells.

**Fig 2:**
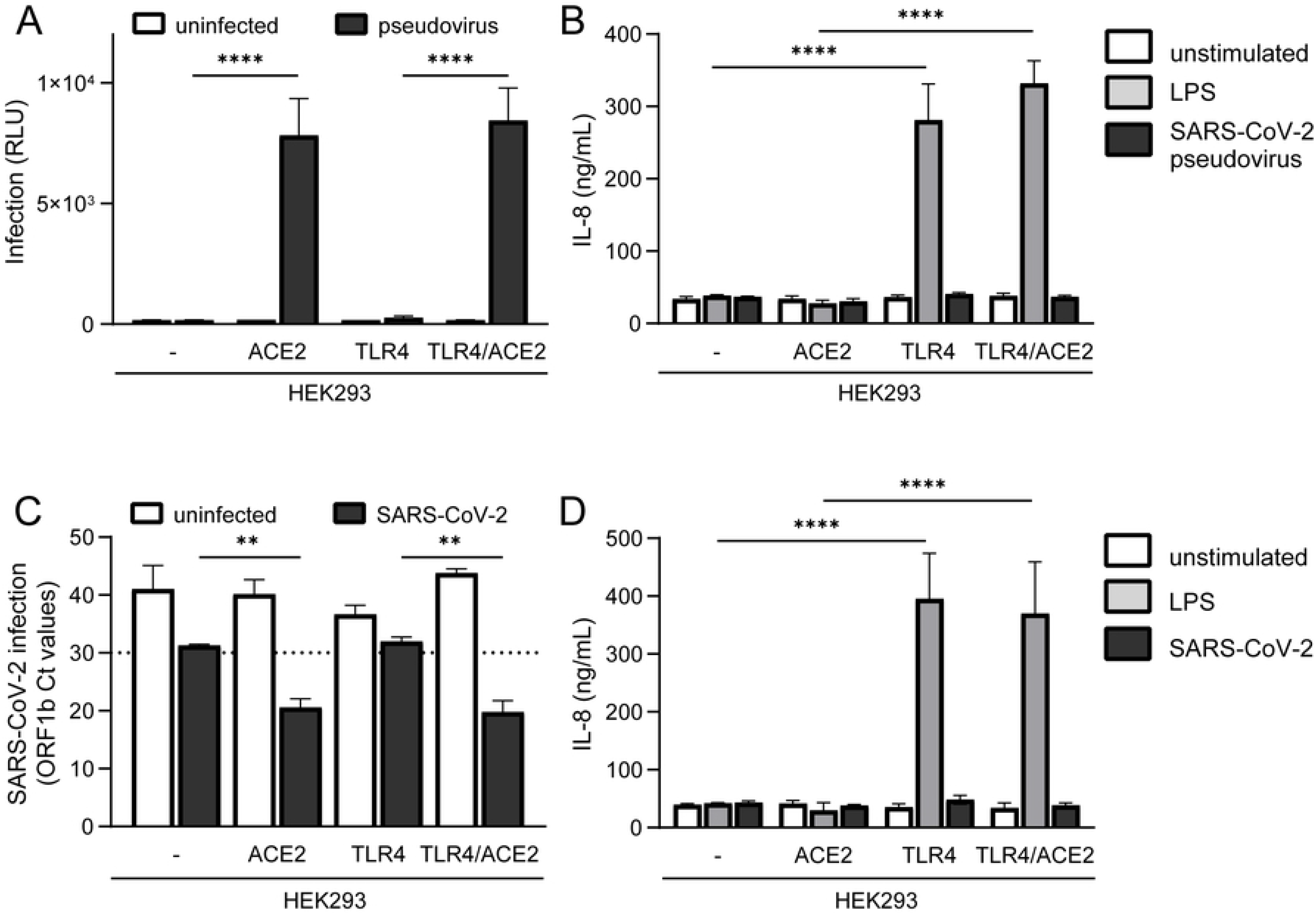
SARS-CoV-2 virus particles do not trigger TLR4. (A-B) ACE2-positive and -negative 293 and 293/TLR4 cells were exposed to SARS-CoV-2 pseudovirus and infection was determined after 3 days by measuring luciferase activity (A), and IL-8 production was measured after 24h by ELISA (B). (C-D) ACE2-positive and -negative 293 and 293/TLR4 cells were exposed to a primary SARS-CoV-2 isolate and infection was determined after 24h by measuring the viral gene ORFb1 expression in supernatant by qPCR (C) and IL-8 production was measured after 24h by ELISA (D). Data show the mean values and SEM. Statistical analysis was performed using two-way ANOVA with Šidák’s (A) or Tukey’s (B-D) multiple comparisons test. (A-D) ****p<0.0001; **p<0.01 (A-B; n=3 in triplicates) (C-D; n=3). RLU = relative light units.

Next we investigated whether SARS-CoV-2 pseudovirus activates TLR4. SARS-CoV-2 pseudovirus did neither induce IL-8 in parental 293 nor in 293/TLR4 cells (Fig 2B). Moreover, ACE2 expression did not induce activation as exposure of ACE2-positive 293 and 293/TLR4 cells to SARS-CoV-2 pseudovirus did not lead to IL-8 production (Fig 2B). These data further support the findings that S protein from SARS-CoV-2 does not trigger TLR4 and also show that ACE2 does not affect TLR4 signaling.

Next, we treated either ACE2-positive or -negative 293 and 293/TLR4 cells with a primary SARS-CoV-2 isolate (hCoV-19/Italy) and determined infection and activation. Infection was determined by measuring virus particles in the supernatant by qPCR. As expected, both 293/ACE2 and 293/TLR4/ACE2 cells were productively infected at similar levels by SARS-CoV-2, in contrast to ACE2-negative 293 and 293/TLR4 cells (cutoff Ct values >30), (Fig 2C). Neither ACE2-positive nor -negative 293 and 293/TLR4 cells expressed any IL-8 upon exposure to the primary SARS-CoV-2 isolate (Fig 2D). These data strongly suggest that TLR4 does not sense infectious SARS-CoV-2 virus particles.

### Infectious SARS-CoV-2 does not activate DCs

Subsequently, we examined whether SARS-CoV-2 pseudovirus induces DC maturation and cytokine production. DCs do not express ACE2 and we have previously shown that SARS-CoV-2 pseudovirus does not infect DCs(23). We investigated the maturation and cytokine production by DCs stimulated with SARS-CoV-2 pseudovirus. Exposure of DCs to SARS-CoV-2 pseudovirus did neither induce expression of costimulatory markers CD80 and CD86 nor maturation marker CD83, in contrast to LPS (Fig 3A-D). Moreover, SARS-CoV-2 pseudovirus did not induce any cytokines, in contrast to LPS (Fig 3E-H). These data indicate that the S protein expressed by SARS-CoV-2 pseudovirus does not activate DCs.

**Fig 3:**
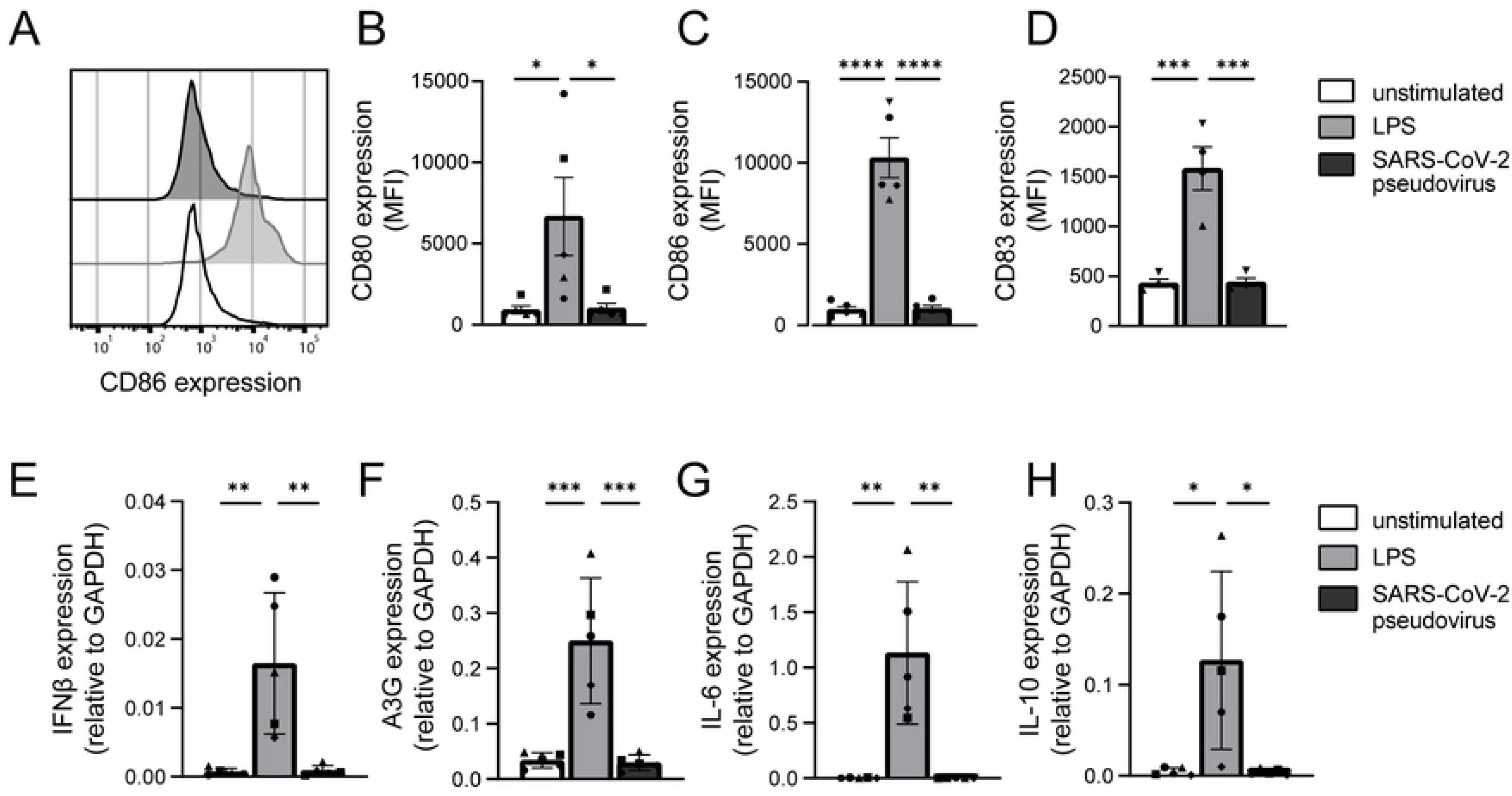
SARS-CoV-2 pseudovirus does not activate dendritic cells. (A-D) Primary DCs were exposed to LPS or SARS-CoV-2 pseudovirus and maturation and cytokine production was determined after 24h and 6h respectively. (A) Representative histogram of CD86 expression. (B-D) Cumulative flow cytometry data of CD80 (B), CD86 (C), and CD83 (D) expression. (E-H) mRNA levels of IFNβ (E), A3G (F), IL-6 (G) and IL-10 (H) were determined with qPCR. Data show the mean values and SEM. Statistical analysis was performed using one-way ANOVA with Tukey’s multiple comparisons test. (B-D) ****p<0.0001; ***p<0.001; *p<0.05 (B-C; n=5) (D; n=4). (E-H) ***p<0.001; **p<0.01; *p<0.05 (n=5). MFI = mean fluorescence intensity.

Next, we exposed DCs to a primary SARS-CoV-2 isolate and determined DC maturation and cytokine production. We have previously shown that DCs do not become infected by primary SARS-CoV-2(23). Exposure of DCs to the primary SARS-CoV-2 isolate did neither induce expression of CD80 CD86, nor CD83, whereas LPS induced expression of CD83 and CD86 (Fig 4A-C).

**Fig 4:**
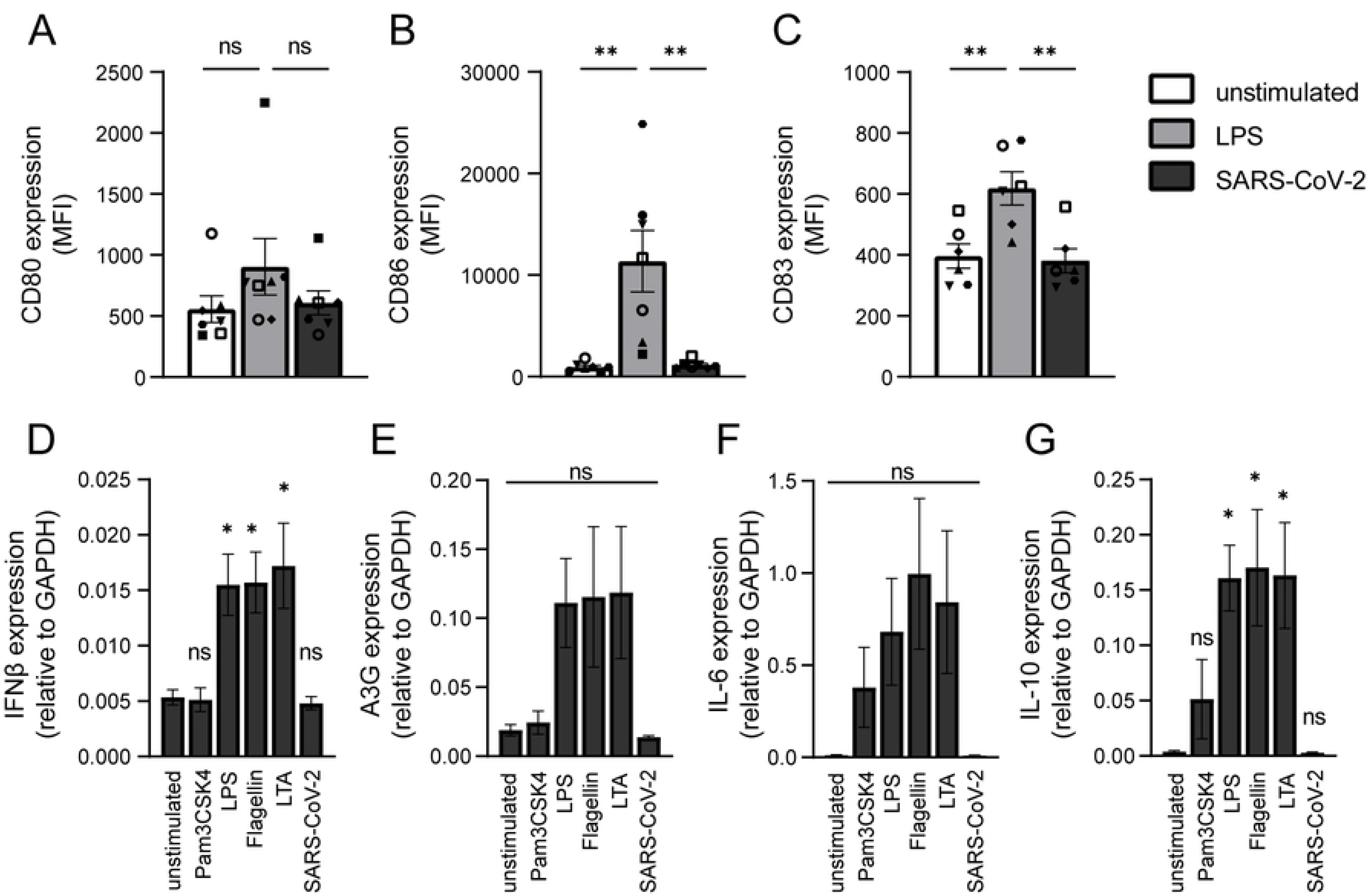
Primary SARS-CoV-2 isolate does not activate dendritic cells. (A-G) Primary DCs were exposed to LPS or primary SARS-CoV-2 isolate and DC maturation was measured after 24h by flow cytometry. Cumulative flow cytometry data of CD80 (A), CD86 (B), and CD83 (C) expression. (D-G) Primary DCs were exposed to different TLR agonists or primary SARS-CoV-2 isolate and mRNA levels of IFNβ (D), A3G (E), IL-6 (F) and IL-10 (G) were determined with qPCR. Data show the mean values and SEM. Statistical analysis was performed using one-way ANOVA with Tukey’s multiple comparisons test. (A-C) **p<0.01; ns = non-significant (A-B; n=7) (C; n=6). (D-G) Data are compared to the unstimulated condition, *p<0.05; ns = non-significant (n=5). MFI = mean fluorescence intensity.

Next we investigated cytokine induction by DCs after exposure to primary SARS-CoV-2 isolate or agonists for extracellular TLRs (TLR1/2, TLR2/6, TLR4, and TLR5). LPS, flagellin and LTA induced type I IFN responses as well as cytokines, whereas Pam3CSK4 only induced cytokines (Fig 4D-G). However, exposure of DCs to the primary SARS-CoV-2 isolate did not lead to induction of type I IFN responses nor cytokines (Fig 4D-G). Therefore, these data strongly indicate that primary SARS-CoV-2 virus particles are not sensed by any extracellular PRRs on DCs such as TLR2, TLR4, and TLR5.

### Ectopic ACE2 expression on DCs results in SARS-CoV-2 infection and immune activation

Next, we investigated whether infection of DCs after ectopic expression of ACE2 with primary SARS-CoV-2 isolate would induce immune responses. DCs do not express ACE2, but transfection with ACE2 plasmid resulted in ACE2 mRNA and surface expression (Fig 5A-C). Next, both mock- and ACE2-transfected DCs were exposed to the primary SARS-CoV-2 isolate for 24h in presence or absence of blocking antibodies against ACE2. ACE2-expressing DCs were infected by SARS-CoV-2 and infection was blocked by antibodies against ACE2 (Fig 5D). Notably, infection of DCs with SARS-CoV-2 induced transcription of IFNβ (Fig 5E) as well as the ISG A3G (Fig 5F). Infection also induced pro-inflammatory cytokine IL-6 (Fig 5G). Both type I IFN responses and IL-6 were abrogated by blocking infection using ACE2 antibodies. Taken together, these data strongly indicate that infection is required to induce cytokine responses by DCs and suggest that intracellular PRRs rather than extracellular TLRs are involved in sensing SARS-CoV-2 and instigating immune responses against SARS-CoV-2.

**Fig 5:**
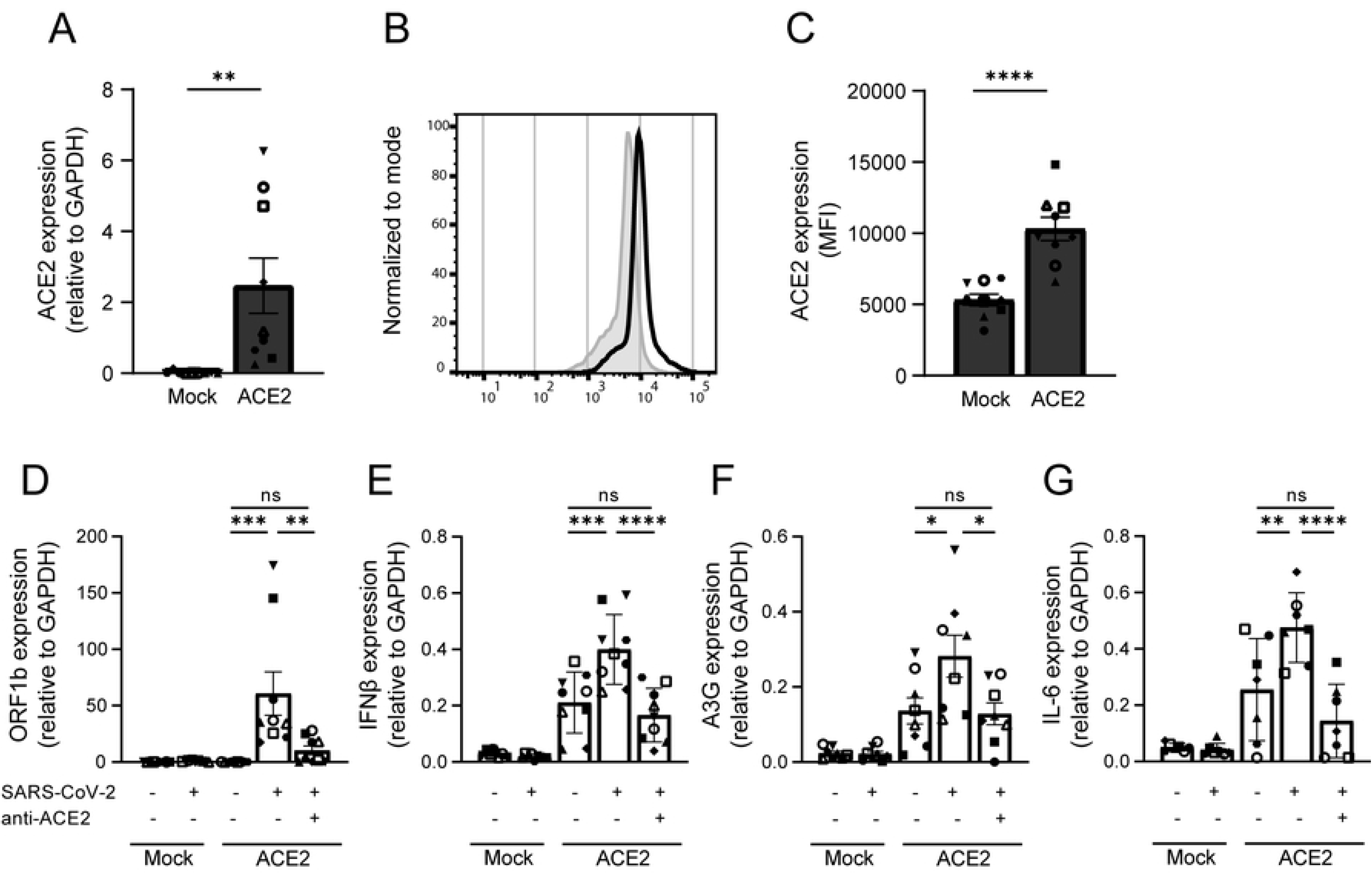
Ectopic expression of ACE2 on DCs results in infection and induction of immune responses. (A-C) Ectopic expression of ACE2 on primary DCs was determined by qPCR and flow cytometry. (A) Cumulative qPCR data of ACE2 expression on DCs. (B) Representative histogram of ACE2 expression on DCs. (C) Cumulative flow cytometry data of ACE2 expression. (D-G) ACE2-positive and -negative DCs were exposed to primary SARS-CoV-2 isolate in presence or absence of blocking antibodies against ACE2. Infection (D) and mRNA levels of IFNβ (E), A3G (F), and IL-6 (G) were determined with qPCR. Data show the mean values and SEM. Statistical analysis was performed using (A, C) unpaired student’s t-test or (D-G) one-way ANOVA with Tukey’s multiple comparisons test. (A, C) ****p<0.0001; **p<0.01 (n=9). (D-G) ****p<0.0001; ***p<0.001; **p<0.01; *p<0.05; ns = non-significant; (D-F; n=9) (G; n=7). MFI = mean fluorescence intensity.

## Discussion

SARS-CoV-2 has established itself as a contagious human respiratory pathogen, which can trigger a robust inflammatory cytokine response(8). However, it remains largely unknown whether innate immune receptors are involved in the onset of immune responses against SARS-CoV-2. TLR4 has been suggested to play a role in sensing SARS-CoV-2 and inducing a strong immune response(13, 14). Here, our data suggest that SARS-CoV-2 by itself is not recognized by TLR4, as neither a TLR4-expressing 293 cell line nor primary DCs were activated by exposure to recombinant S protein, SARS-CoV-2 pseudovirus or primary SARS-CoV-2 virus particles. Ectopic expression of ACE2 on primary DCs allowed infection with primary SARS-CoV-2. Notably, productive infection of ACE2-positive DCs induced type I IFN and cytokine responses, which was abrogated by blocking ACE2. Our data therefore suggest that SARS-CoV-2 virus particles are not sensed by extracellular TLRs, including TLR4, but that infection via ACE2 is required.

Other studies have reported that S protein triggered TLR4, and also TLR2 and TLR6 are suggested to interact with SARS-CoV-2 S protein to induce pro-inflammatory responses(13, 14, 17–19, 24). However, neither a TLR4-expressing 293 cell line nor primary DCs were activated by recombinant S proteins. As it is possible that contaminations during the purification process of recombinant proteins might induce activation, we also investigated immune activation by SARS-CoV-2 pseudovirus and infectious primary SARS-CoV-2. However, neither TLR4-expressing 293 cells nor primary DCs were activated by pseudovirus or a primary isolate of SARS-CoV-2 as measured by cytokine production and DC maturation. Therefore, our data strongly suggest that S protein expressed by SARS-CoV-2 does not trigger TLR4. Since monocyte-derived DCs do not express ACE2, they are not infected by SARS-CoV-2 and therefore exposure to primary SARS-CoV-2 will only allow sensing by extracellular PRRs that are expressed by DCs. Therefore, our data also imply that extracellular transmembrane TLRs do not sense SARS-CoV-2 virus particles.

Notably, ectopic expression of ACE2 on monocyte-derived DCs leads to infection and the production of cytokines, suggesting that replication of SARS-CoV-2 triggers cytosolic sensors. Indeed, studies suggest that intracellular viral sensors such as RIG-I or MDA5 are involved in SARS-CoV-2 infection(25–27).

Interestingly, our data suggest that infection of immune cells and thus antigen presenting cells (APCs) is essential to induction of immunity. Therefore, it is important to identify ACE2-positive DC subsets and macrophages, since these APCs should be sensitive to infection and are therefore paramount in initiating adaptive immunity. In the absence of DC infection, epithelial cell infection and subsequent inflammation and tissue damage might account for initial immune activation and inflammation, and subsequent release of PAMPs and DAMPs might activate DCs(28). It remains unclear whether these secondary signals are able to correctly instruct DCs and this might underlie the strong inflammatory responses observed during COVID-19. Our finding that SARS-CoV-2 is not recognized by TLR4 might therefore be an escape mechanism leading to inefficient DC activation and subsequent aberrant inflammatory responses.

It has been suggested that worsening of disease in COVID-19 patients coincides with the activation of the adaptive immune response, 1-2 weeks after infection(8). Since DCs have a bridging function to activate the adaptive immune response, it is important to study DCs in the context of COVID-19. Our research suggests that ACE2-negative DCs are not properly activated by infectious SARS-CoV-2. Moreover, our data suggest that SARS-CoV-2 is able to escape from extracellular TLRs that are one of the most important PRR families crucial for induction of innate and adaptive immunity, and further research will show whether the lack of TLR activation underlies observed inflammation during COVID-19.

## Materials and methods

### Cell lines

The Simian kidney cell line VeroE6 (ATCC® CRL-1586™) was maintained in CO_2_ independent medium (Gibco Life Technologies, Gaithersburg, Md.) supplemented with 10% fetal calf serum (FCS), 2mM L-glutamine and penicillin/streptomycin. Culture was maintained at 37°C without CO_2_.

Human embryonic kidney cells (HEK293) were maintained in IMDM (Gibco) supplemented with 10% FCS and 1% penicillin/streptomycin (Invitrogen). HEK293 cells stably transfected with TLR4 cDNA (HEK/TLR4) were a kind gift from D. T. Golenbock(15). HEK293 and HEK/TLR4 cells were transiently transfected with pcDNA3.1(-)hACE2 (Addgene plasmid #1786) to generate HEK/ACE2 or HEK/TLR4/ACE2 cell lines. Transfection was performed using Lipofectamine LTX and PLUS reagent (Invitrogen) according to the manufacturer’s protocol. After 24h, cells were split and seeded into flat-bottom 96-well plates (Corning) and left to attach for 24h, before performing further experiments. Cultures were maintained at 37°C and 5% CO_2_. Before infection with the SARS-CoV-2 isolate (described below), media was exchanged for CO_2_-independent media, since infection with a SARS-CoV-2 primary isolate occurs under CO_2_ negative conditions. Human ACE2-expressing cell lines were analyzed for ACE2 expression via quantitative real-time PCR.

### Primary cells

This study was performed in accordance with the ethical principles set out in the declaration of Helsinki and was approved by the institutional review board of the Amsterdam University Medical Centers, location AMC Medical Ethics Committee and the Ethics Advisory Body of Sanquin Blood Supply Foundation (Amsterdam, Netherlands). Human CD14+ monocytes were isolated from the blood from healthy volunteer donors (Sanquin blood bank) and subsequently differentiated into monocyte-derived dendritic cells (DCs). The isolation from buffy coats was done by density gradient centrifugation on Lymphoprep (Nycomed) and Percoll (Pharmacia). After separation by Percoll, the isolated monocytes were cultured in RPMI 1640 (Gibco) supplemented with 10% FCS, 2mM L-glutamin (Invitrogen) and 10 U/mL penicillin and 100 ug/mL streptomycin, containing the cytokines IL-4 (500 U/mL) and GM-CSF (800 U/mL) (both Gibco) for differentiation into DCs. After 4 days of differentiation, DCs were seeded at 1×10^6^ /mL in a 96-well plate (Greiner), and after 2 days of recovery, DCs were stimulated or infected as described below.

Alternatively, monocyte-derived DCs that were transfected with hACE2 were seeded at 0.5×10^6^ cells/mL in a 6-well plate and transfection was performed with Lipofectamine LTX and PLUS reagents (Invitrogen) according to the manufacturer’s instructions for primary cells. After 24h, cells were seeded at 1×10^6^/mL in a 96-well plate and after 24h of recovery, they were infected with primary SARS-CoV-2 isolate.

### SARS-CoV-2 pseudovirus production

For production of single-round infection viruses, human embryonic kidney 293T/17 cells (ATCC, CRL-11268) were co-transfected with an adjusted HIV-1 backbone plasmid (pNL4-3.Luc.R-S-) containing previously described stabilizing mutations in the capsid protein (PMID: 12547912) and firefly luciferase in the *nef* open reading frame (1.35ug) and pSARS-CoV-2 expressing SARS-CoV-2 S protein (0.6ug) (GenBank; MN908947.3)(22). Transfection was performed in 293T/17 cells using genejuice (Novagen, USA) transfection kit according to manufacturer’s protocol. At day 3 or day 4, pseudotyped SARS-CoV-2 virus particles were harvested and filtered over a 0.45 μm nitrocellulose membrane (SartoriusStedim, Gottingen, Germany). SARS-CoV-2 pseudovirus productions were quantified by p24 ELISA (Perkin Elmer Life Sciences).

### SARS-CoV-2 (primary isolate) virus production

The following reagent was obtained from Dr. Maria R. Capobianchi through BEI Resources, NIAID, NIH: SARS-Related Coronavirus 2, Isolate Italy-INMI1, NR-52284, originally isolated January 2020 in Rome, Italy. VeroE6 cells (ATCC® CRL-1586™) were inoculated with the SARS-CoV-2 isolate and used for reproduction of virus stocks. CPE formation was closely monitored and virus supernatant was harvested after 48 hours. Tissue culture infectious dose (TCID50) was determined on VeroE6 cells by MTT assay 48 hours after infection. Loss of MTT staining as determined by spectrometer is indicative of cell death.

### Stimulation and infection

HEK293 and transfected derivatives were left unstimulated or stimulated for 24h with 10 ng/mL lipopolysaccharide (LPS) from Salmonella (Sigma), 10 ug/mL isolated S protein, 10 ug/mL S nanoparticle, or with pseudotyped or authentic SARS-CoV-2, as specified below. DCs were left unstimulated, or stimulated with 10 ug/ml Pam3CSK4 (Invivogen), 10 ng/mL LPS from *Salmonella typhosa* (Sigma), 10 ug/mL flagellin from *Salmonella typhimurium* (Invivogen), 10 ug/mL lipoteichoic acid (LTA) from *Staphylococcus aureus* (Invivogen), pseudotyped virus or SARS-CoV-2. Blocking of ACE2 was performed with 8 ug/mL anti-ACE2 (R&D systems) for 30 min at 37°C before adding stimuli. Monocyte-derived DCs do not express ACE2 and are therefore not infected. Therefore, pseudovirus stimulation was performed for 6h, after which the cells were lysed for mRNA analysis of cytokine production. DCs ectopically expressing ACE2 were stimulated for 24h with virus before the cells were lysed for mRNA analysis of cytokine production. Also, cells were stimulated for 24h and fixed for 30 min with 4% paraformaldehyde, after which the expression of maturation markers was assessed with flow cytometry.

For the pseudovirus infection assays, HEK293 or 293/TLR4 cell lines and DCs were exposed to 95ng/mL and 191.05ng/mL of SARS-CoV-2 pseudovirus, respectively. Viral protein production was quantified after 3 days at 37°C by measuring luciferase reporter activity. Luciferase activity was measured using the Luciferase assay system (Promega, USA) according to manufacturer’s instructions.

For the primary SARS-CoV-2 infection assays, HEK293 or HEK/TLR4 cell lines and DCs were exposed to the SARS-CoV-2 isolate (hCoV-19/Italy) at different TCIDs (100 and 1000; MOI 0.0028-0.028) for 24 hours at 37°C. After 24 hours, cell supernatant was taken and DCs were lysed for isolation of viral RNA. Also, the HEK293/ACE2 and HEK/TLR4/ACE2 cell lines were exposed to the SARS-CoV-2 isolate (hCoV-19/Italy) at TCID 100 (MOI 0.0028) for 24 hours at 37°C. After 24 hours, the cells were washed 3 times and new media was added. After 48h, cell supernatant was harvested and the cells were lysed to investigate productive infection.

### RNA isolation and quantitative real-time PCR

Cells exposed to SARS-CoV-2 pseudovirus were lysed and mRNA was isolated with the mRNA Catcher™ PLUS Purification Kit (ThermoFisher). Subsequently, cDNA was synthesized with a reverse-transcriptase kit (Promega). RNA of cells exposed to SARS-CoV-2 WT was isolated with the QIAamp Viral RNA Mini Kit (Qiagen) according to the manufacturer’s protocol. cDNA was synthesized with the M-MLV reverse-transcriptase kit (Promega) and diluted 1 in 5 before further application. PCR amplification was performed in the presence of SYBR green (ThermoFisher) in a 7500 Fast Realtime PCR System (ABI). Specific primers were designed with Primer Express 2.0 (Applied Biosystems). The ORF1b primers used were as described before(29). The normalized amount of target mRNA was calculated from the Ct values obtained for both target and household mRNA with the equation Nt = 2^Ct(GAPDH)-Ct(target)^. The following primers were used:

GAPDH: F_CCATGTTCGTCATGGGTGTG; R_GGTGCTAAGCAGTTGGTGGTG; TLR4:
F_CTGCAATGGATCAAGGACCAG; R_CCATTCGTTCAACTTCCACCA; ACE2:
F_GGACCCAGGAAATGTTCAGA; R_ GGCTGCAGAAAGTGACATGA; ORF1b:
F_TGGGGTTTTACAGGTAACCT; R_AACACGCTTAACAAAGCACTC; IL-8: F_TGAGAGTGGACCACACTGCG;
R_TCTCCACAACCCTCTGCACC; IFNB: F_ACAGACTTACAGGTTACCTCCGAAAC;
R_CATCTGCTGGTTGAAGAATGCTT; APOBEC3G: F_TTGAGCCTTGGAATAATCTGCC;
R_TCGAGTGTCTGAGAATCTCCCC; IL-6: F_TGCAATAACCACCCCTGACC;
R_TGCGCAGAATGAGATGAGTTG; IL-10: F_GAGGCTACGGCGCTGTCAT; R_CCACGGCCTTGCTCTTGTT

### ELISA

Cell supernatants were harvested after 24h of stimulation and secretion of IL-8 was measured by ELISA (eBiosciences) according to the manufacturer’s instructions. OD450 nm values were measured using a BioTek Synergy HT. Supernatant containing SARS-CoV-2 pseudovirus was inactivated with 0.1% triton and supernatant containing SARS-CoV-2 was inactivated with 1% triton before performing ELISA.

### Flow cytometry

For cell surface staining, cells were incubated in 0.5% PBS-BSA (phosphate-buffered saline containing 0.5% bovine serum albumin (BSA; Sigma-Aldrich)) containing antibodies for 30 min at 4°C. Single-cell measurements were performed on a FACS Canto flow cytometer (BD Biosciences) and FlowJo V10 software (TreeStar) was used to analyze the data. The antibody clones used are: CD86 (2331 (FUN-1), BD Pharmingen), CD80 (L307.4, BD Pharmingen), CD83 (HB15e, BD Pharmingen), ACE2 (AF933, R&D systems), goat-IgG (AB-2535864, ThermoFisher Scientific), donkey-anti-goat (A-21447, ThermoFisher Scientific). For each experiment, live cells were gated on FSC and SSC and analyzed further with the markers mentioned.

### Statistics

Graphpad Prism v8 (GraphPad Software) was used to generate all graphs and for statistical analyses. Statistics were performed using a Student’s *t* test for pairwise comparisons. Multiple comparisons within groups were performed using an RM one-way analysis of variance (ANOVA) with a Tukey’s multiple comparisons test, or two-way ANOVA with a Tukey’s or Šidák’s multiple comparisons test where indicated. *p* < 0.05 were considered statistically significant.

## Author Contributions

LEHvdD and MBJ designed experiments; LEHvdD, MBJ, JE, and JLvH performed the experiments; PJMB, MB, ACvN, NAK, MJvG and RWS contributed essential research materials and scientific input. LEHvdD, MBJ and TBHG analyzed and interpreted data; LEHvdD, MBJ and TBHG wrote the manuscript with input from all listed authors. TBHG supervised all aspects of this study.

## Data availability

The data generated during this study are available from the corresponding author on reasonable request.

